# Light induced expression of gRNA allows for optogenetic gene editing of T lymphocytes in vivo

**DOI:** 10.1101/2023.11.09.566272

**Authors:** Diego Velasquez Pulgarin, Nathalie Pelo, Lin Ferrandiz, Tilen Trselic, William A Nyberg, Gary Bowlin, Alexander Espinosa

## Abstract

There is currently a lack of tools capable of perturbing genes in both a precise and spatiotemporal fashion. CRISPR’s ease of use and flexibility, coupled with light’s unparalleled spatiotemporal resolution deliverable from a controllable source, makes optogenetic CRISPR a well-suited solution for precise spatiotemporal gene perturbations. Here we present a new optogenetic CRISPR tool, BLU-VIPR, that diverges from prevailing split-Cas design strategies and instead focuses on optogenetic regulation of gRNA production. This simplifies spatiotemporal gene perturbation and works in vivo with cells previously intractable to optogenetic gene editing. We engineered BLU-VIPR around a new potent blue-light activated transcription factor and ribozyme-flanked gRNA. The BLU-VIPR design is genetically encoded and ensures precise excision of multiple gRNAs from a single mRNA transcript, allowing for optogenetic gene editing in T lymphocytes in vivo.

## MAIN

Clustered regularly interspaced short palindromic repeats (CRISPR) and their associated RNA-guided nucleases (Cas) have revolutionized genome engineering, and new systems and functionalities are being developed at a rapid pace. However, to harness the full power of CRISPR, it will be invaluable to establish spatiotemporal control of CRISPR systems in vivo. Since optogenetics is the preferred method for precise spatiotemporal control, optogenetic CRISPR has emerged as a promising method for spatiotemporal gene editing. Most current optogenetic CRISPR systems are based on light-induced dimerization of split-Cas^1^, dimerization of Cas with effectors^2^, or dissociation of anti-CRISPR proteins^3^. Although powerful, these systems have several disadvantages, including the requirement for cumbersome re-engineering to expand their use to additional Cas types and effectors, and their large sizes preventing delivery to primary cells using viral transduction. Alternatively, photocaged RNA can be used for optogenetic CRISPR, but their limited half-life and short persistence *in vivo* reduces their usefulness^4,5^. To circumvent these limitations, we have generated a new optogenetic platform, Blue Light-inducible Universal VPR-Improved Production of RGRs (BLU-VIPR), allowing for simultaneous light-induced expression of genetically encoded single-guide RNAs (gRNA) and proteins. BLU-VIPR can be combined with multiple Cas types and effectors and be delivered by virus for optogenetic gene editing in vivo.

## RESULTS

### Simultaneous induction of protein and gRNA from a blue-light induced pol II promoter

To enable light-induced expression of gRNAs, we took advantage of the blue-light activated protein EL222^6^. Upon exposure to blue light (470 nm), EL222 homodimerizes and binds to its response elements in the C120 promoter^6^. By fusing EL222 to a transcriptional activator it is therefore possible to activate RNA polymerase II (RNAPII)-mediated transcription from the C120 promoter by exposure to blue light (Extended Data Fig. 1a). To ensure strong expression of gRNA after light exposure, we engineered a potent blue-light induced transcription factor by fusing EL222 to the synthetic transcriptional activator VP64-p65-Rta (VPR)^7^. To enable the release of a functional gRNA from the resulting transcript, we flanked the gRNA with self-cleaving hammerhead (HH) and Hepatitis delta virus (HDV) ribozymes in a ribozyme-gRNA-ribozyme (RGR) design^8–10^ (Fig. 1a). These ribozymes precisely excise the gRNA from the transcript and allow for simultaneous production of functional gRNAs and proteins from the same transcript. We then generated a VPR-EL222 expressing construct with both RGR and mCherry under the control of the C120 promoter (Fig. 1b). To test the construct, we used light emitting diode (LED) transilluminators for controllable delivery of blue light (470 nm) to transfected HEK293T cells. Indeed, VPR-EL222 induced high levels of mCherry only after exposure to blue light, with no observable leakiness when kept in dark conditions (Fig. 1c, Extended Data Fig. 1b). In contrast, a fusion of EL222 to the commonly used transcriptional activator VP16 (VP16-EL222) was unable to induce mCherry (Fig. 1c). The robust VPR-EL222-based system for light-induced expression of gRNA was designated BLU-VIPR (Extended Data Fig. 2a, b).

**Figure 1.**
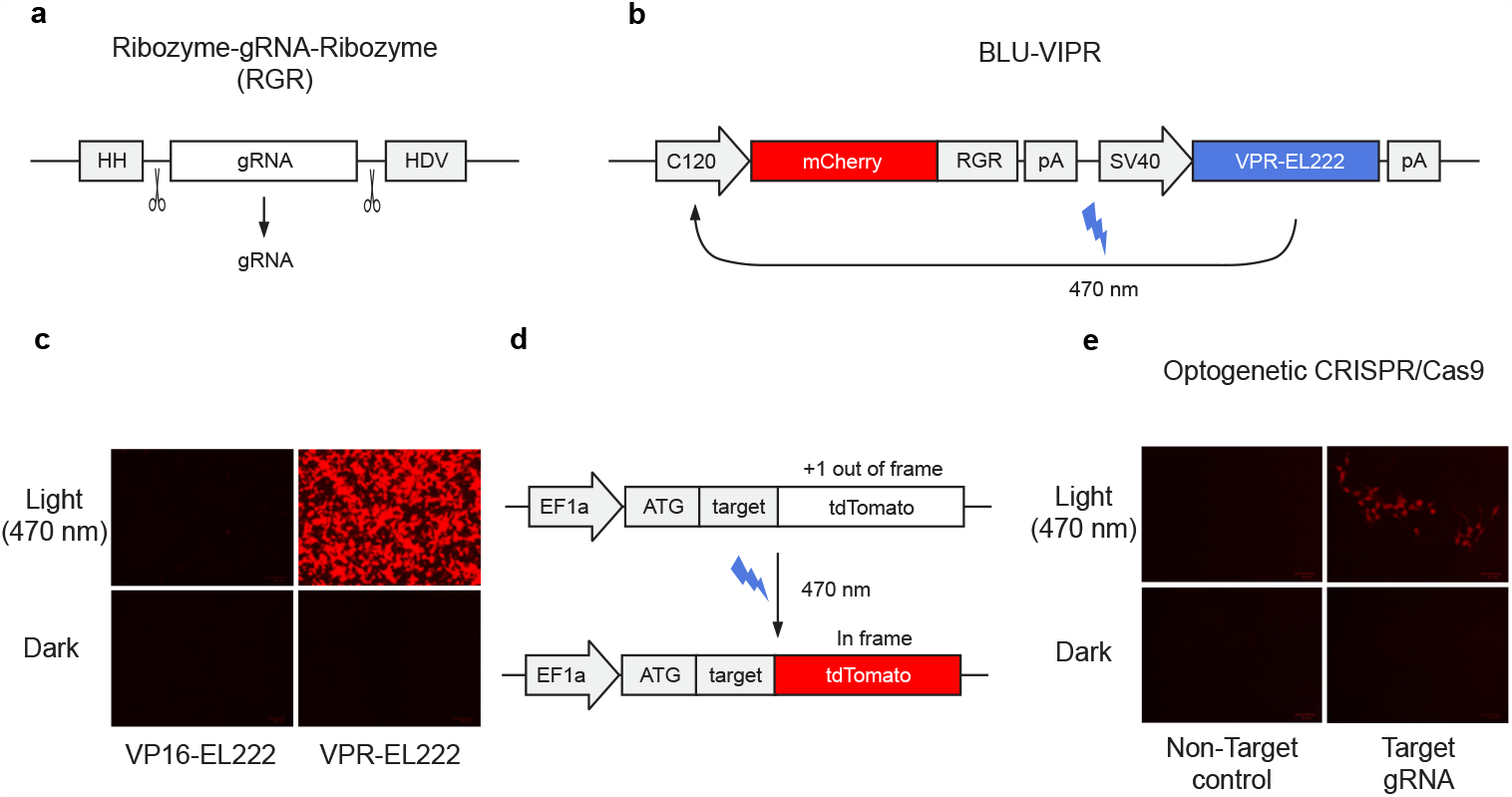
Optogenetic CRISPR using light-induced expression of gRNA. **a**, The ribozyme-gRNA-ribozyme (RGR) design consists of a hammerhead (HH) ribozyme followed by the gRNA and a Hepatitis delta virus (HDV) ribozyme. After precise self-cleavage of the ribozymes a functional gRNA is released. **b**, Design of construct for VPR-EL222 (BLU-VIPR) dependent activation of C120 promoter transcription, allowing for simultaneous expression of mCherry and gRNA after exposure to blue light (470 nm). **c**, Comparison of mCherry reporter expression in HEK293T cells after transfection with VPR-EL222 (BLU-VIPR) or VP16-EL222 constructs followed by exposure to blue light for 24 hours. The experiment was performed three times and one representative image is shown. **d**, Out of frame Cas9 reporter where the expression of tdTomato is restored after Cas9 mediated indels. **e**, HEK293T cells were transfected with BLU-VIPR (without mCherry) containing a gRNA targeting the out of frame sequence in tdTomato. After 48 hours of light exposure, the cells were cultured for an additional 72 hours before detection of tdTomato demonstrating successful optogenetic induction of indels by Cas9. The experiment was performed three times and one representative image is shown.

To establish if BLU-VIPR could be used for optogenetic induction of Cas9 mediated dsDNA breaks, we generated Cas9 reporter cells containing an out of frame (+1 bp) tdTomato under control of the EF1α promoter^11^ (Fig. 1d). Cas9 reporter cells were then co-transfected with a Cas9 plasmid and a BLU-VIPR plasmid containing RGRs with a gRNA targeting the frameshifted sequence of tdTomato, or a non-targeting control gRNA (we removed mCherry from the BLU-VIPR construct to avoid interference with tdTomato). Indeed, we could verify that exposure to blue light restored the reading frame of tdTomato indicating successful optogenetic control of Cas9 (Fig. 1e). In summary, we engineered a system, BLU-VIPR, for potent induction of functional gRNA and protein from the same transcript upon exposure to blue light.

### Multiplexed and orthogonal optogenetic CRISPR activation

Since the RGR design of BLU-VIPR allows for the expression of multiple gRNAs from the same light-induced RNA transcript we reasoned that it should enable multiplexed and orthogonal optogenetic CRISPR. To demonstrate this, we used the BLU-VIPR system for multiplexed and orthogonal CRISPR activation (CRISPRa) based on dCas9-VPR and dCas12a-VPR. We achieved this by inserting two RGRs into the BLU-VIPR construct. The first RGR contained a gRNA for dCas12a-VPR-dependent activation of *PDGFB* (Platelet Derived Growth Factor subunit B), and the second RGR contained a gRNA for dCas9-VPR-dependent activation of *BMP2* (Bone morphogenetic protein 2) (Fig. 2a). This multiplexed RGR design thus allowed for simultaneous expression of two gRNAs from the same transcript. We then co-transfected the multiplexed BLU-VIPR construct together with dCas12a-VPR and dCas9-VPR and measured the induction of *PDGFB* and *BMP2* expression after exposure to blue light. Indeed, the multiplexed BLU-VIPR was able to achieve robust optogenetic activation of both *PDGFB* and *BMP2* expression (Fig. 2b). Importantly, the multiplexed RGR design was truly orthogonal since the *PDFGB*-specific gRNA was unable to activate *PDFGB* transcription with dCas9-VPR, and conversely, the *BMP2*-specific gRNA was unable to activate *BMP2* transcription with dCas12a-VPR.

**Figure 2.**
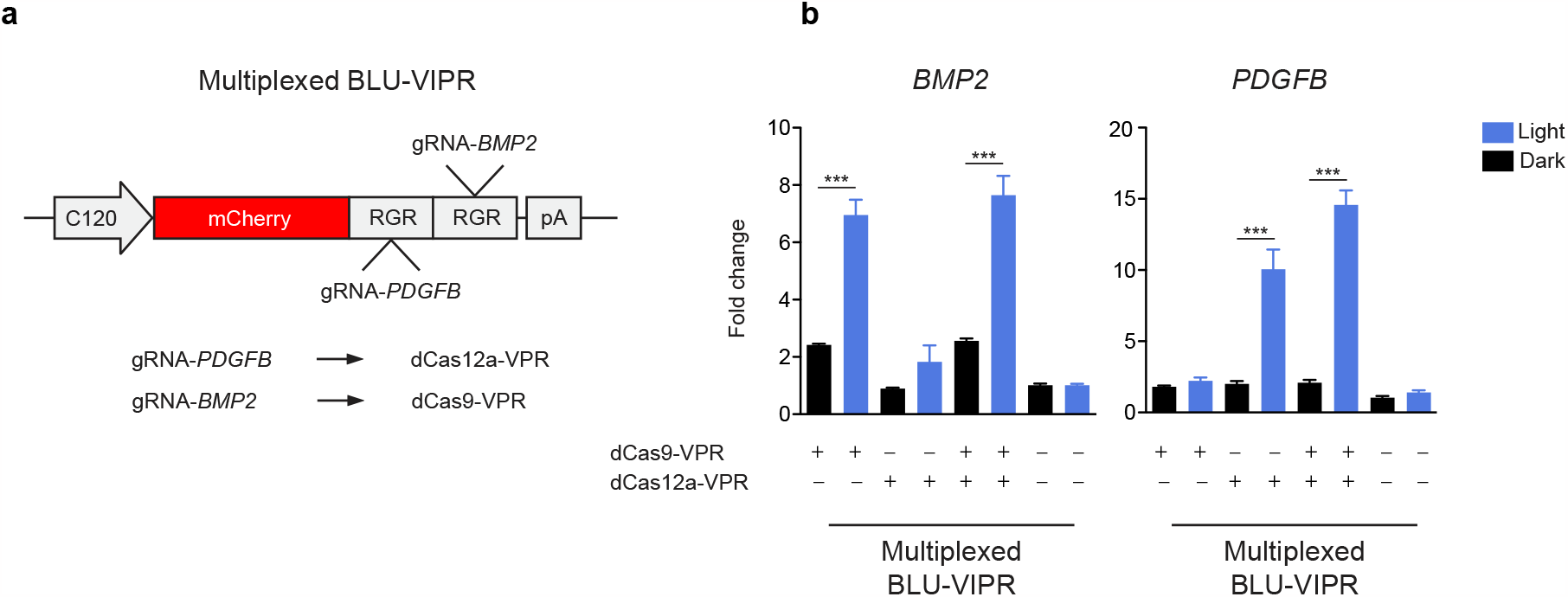
Multiplexed and orthogonal optogenetic CRISPR activation. **a**, Ribozyme-gRNA-ribozyme (RGR) design for multiplexed activation of dCas12a-VPR and dCas9-VPR. **b**, Multiplexed, orthogonal optogenetic dCas12a-VPR and Cas9-VPR to activate the endogenous genes *PDGFB* and *BMP2*. HEK293T cells were transfected with BLU-VIPR-multiplexed, and either dCas12a-VPR, dCas9-VPR, or both. *PDGFB* and *BMP2* expression was assayed after 48 hours of light stimulation, normalized to *HPRT1* expression and fold change is relative to untransfected samples kept under dark conditions. The experiment was performed with n=5. One-way ANOVA with Tukey post hoc test was used to identify statistical significance (***p<0.001).

We also compared BLU-VIPR to LACE, a previous optogenetic CRISPRa system based on split-Cas^2^. In contrast to LACE, BLU-VIPR was able to strongly induce *BMP2* with a single gRNA after light exposure (Extended Data Fig. 2c). Altogether, these results demonstrate that BLU-VIPR is a VPR-EL222-based gRNA expression system that allows for multiplexed and orthogonal optogenetic activation of dCas9-VPR and dCas12a-VPR.

### Optogenetic C-to-T and A-to-G base editing

The ability to achieve optogenetic control of both Cas9 and Cas12 demonstrated that BLU-VIPR is compatible with different types of Cas proteins. We therefore asked if BLU-VIPR also could achieve control of Cas variants previously intractable for optogenetic CRISPR. To answer this, we tested if BLU-VIPR could induce C-to-T and A-to-G base editing after light exposure. We first generated C-to-T and A-to-G base editing reporter cells (Extended Data Fig 3a, b), and then co-transfected them with Target-AID^12^ (C-to-T base editor), or ABEmax^13^ (A-to-G base editor), together with BLU-VIPR plasmids containing gRNAs specific for the target sites in the respective reporters. After exposure to blue light, analysis by flow cytometry revealed populations of mCherry^+^ EGFP^+^ cells indicating activation of BLU-VIPR (mCherry^+^) and successful optogenetic C-to-T or A-to-G base editing (EGFP^+^) (Fig. 3a, b). To test if the BLU-VIPR system also was capable of optogenetic base editing of endogenous genes, we transfected 293FT cells with Target-AID and BLU-VIPR containing a *NEAT1* (Nuclear Enriched Abundant Transcript 1) specific gRNA (Fig. 3c). After exposure to blue light, we sorted mCherry^+^ cells by flow cytometry and sequenced the targeted region in *NEAT1* (Fig. 3d). We could demonstrate successful optogenetic C-to-T base editing in the editing window using Sanger sequencing and identification of base edits with EditR^14^ (Fig. 3e). In summary, we demonstrated successful optogenetic induction of C-to-T and A-to-G base editing, including the successful optogenetic C-to-T base editing of an endogenous gene. To our knowledge, this is the first demonstration of successful optogenetic base editing.

**Figure 3.**
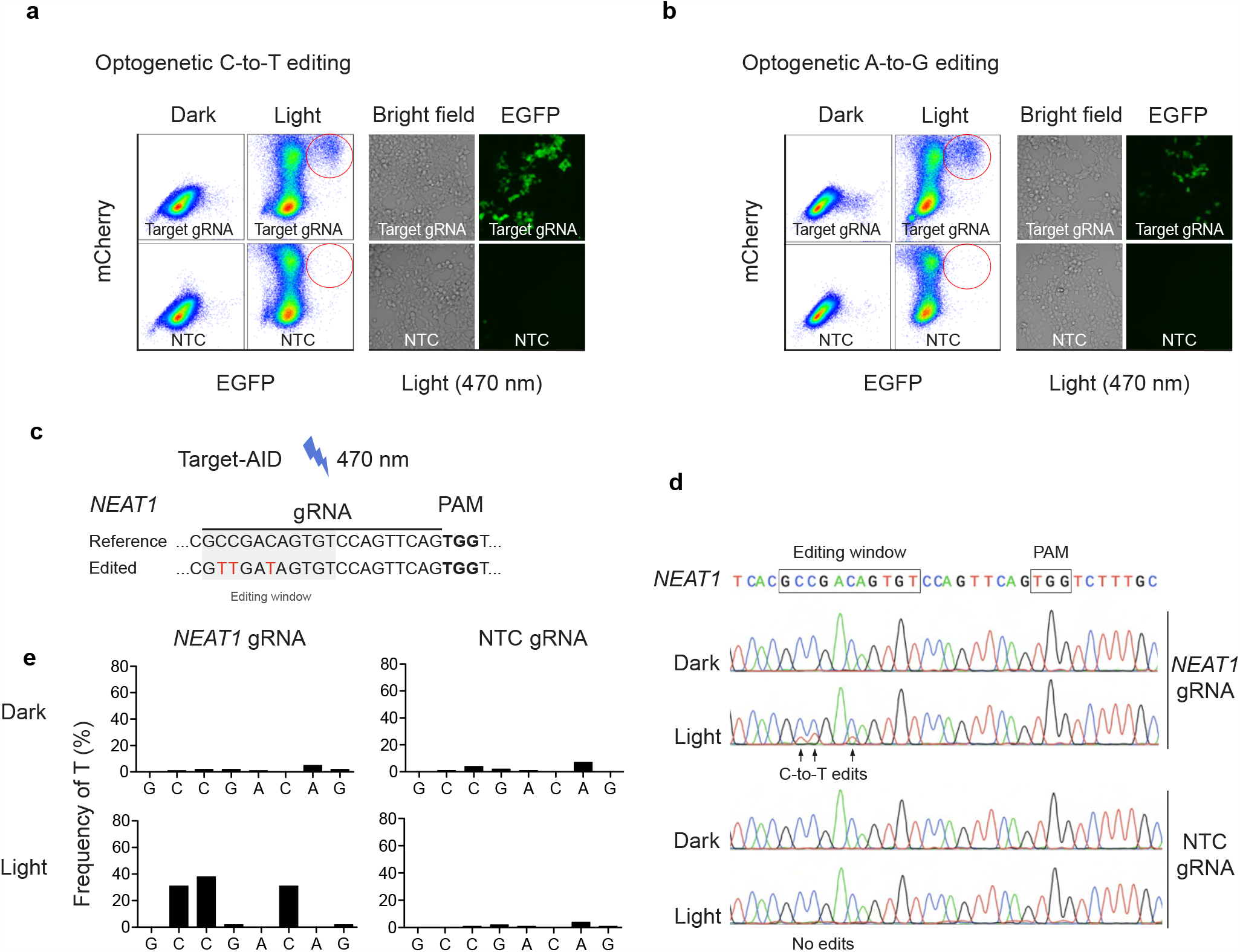
Optogenetic C-to-T and A-to-G base editing. **a**, C-to-T reporter HEK293T cells, expressing EGFP upon successful C-to-T base editing, were transfected with Target-AID and BLU-VIPR containing a gRNA specific for the relevant target sequence in EGFP, or a non-targeting gRNA. Reporter cells were then illuminated, or kept under dark conditions, for 48 hours and analyzed by flow cytometry. Detection of mCherry^+^ EGFP^+^ cells is evidence of successful optogenetic C-to-T base editing. Fluorescence microscopy confirmed presence of EGFP^+^ cells after optogenetic C-to-T base editing in reporter cells. The experiment was repeated three times and one representative result is shown. **b**, A-to-G reporter HEK293T cells, expressing EGFP upon successful A-to-G base editing, were transfected with ABEmax and BLU-VIPR containing a gRNA specific for the relevant target sequence in EGFP, or a non-targeting gRNA. Reporter cells were then illuminated, or kept under dark conditions, for 48 hours and analyzed by flow cytometry. Detection of mCherry^+^ EGFP^+^ cells is evidence of successful optogenetic A-to-G base editing. The experiment was repeated three times and one representative result is shown. **c**, Design of gRNA for optogenetic C-to-T base editing of *NEAT1*. **d**, 293FT cells were transfected with Target-AID and BLU-VIPR plasmids containing non-targeting control (NTC) or *NEAT1*-specific gRNA followed by exposure to blue light. Frequencies of C-to-T edits in the editing window were determined by Sanger sequencing. Representative data from two experiments is shown. **e**, Frequency of C-to-T edits in *NEAT1* after exposure to light.

### Optogenetic gene editing in primary T lymphocytes

Optogenetic gene editing in hard-to-transfect primary cells is challenging due to the necessity of delivering large Cas-based optogenetic machineries into cells^15^. In contrast, the small size of the BLU-VIPR system allows it to be introduced to primary cells using efficient viral delivery. To demonstrate that BLU-VIPR can be used for optogenetic CRISPR gene editing in primary cells, we transduced Cas9 transgenic primary mouse T lymphocytes with a retrovirus containing BLU-VIPR. First, we generated murine stem cell virus (MSCV) constructs for retroviral delivery of BLU-VIPR (MSCV-BLU-VIPR) containing a gRNA specific for *Thy1* (Thy1.2) or an NTC gRNA (Fig. 4a). We then isolated splenic T lymphocytes from Cas9 transgenic mice^16^ followed by transduction with MSCV-BLU-VIPR retrovirus and exposure to pulsed blue light for 48 hours (Fig. 4b). After light exposure, the T lymphocytes were left in the dark for 72 hours before analysis of Thy1.2 expression by flow cytometry (gating strategy in Extended Data Fig. 4a). Indeed, Thy1.2 knockout T lymphocytes were only detected after transduction with Thy1.2 specific MSCV-BLU-VIPR and exposure to blue light (Fig. 4c). We did not detect loss of Thy1.2 either in T lymphocytes that were transduced with Thy1.2 specific MSCV-BLU-VIPR virus and kept in the dark, or in T lymphocytes transduced with NTC control MSCV-BLU-VIPR virus and exposed to blue light. Altogether, these results demonstrate that BLU-VIPR enables optogenetic gene editing in primary T lymphocytes. To our knowledge this makes BLU-VIPR the first optogenetic CRISPR system to allow for optogenetic gene editing in primary lymphocytes.

**Figure 4.**
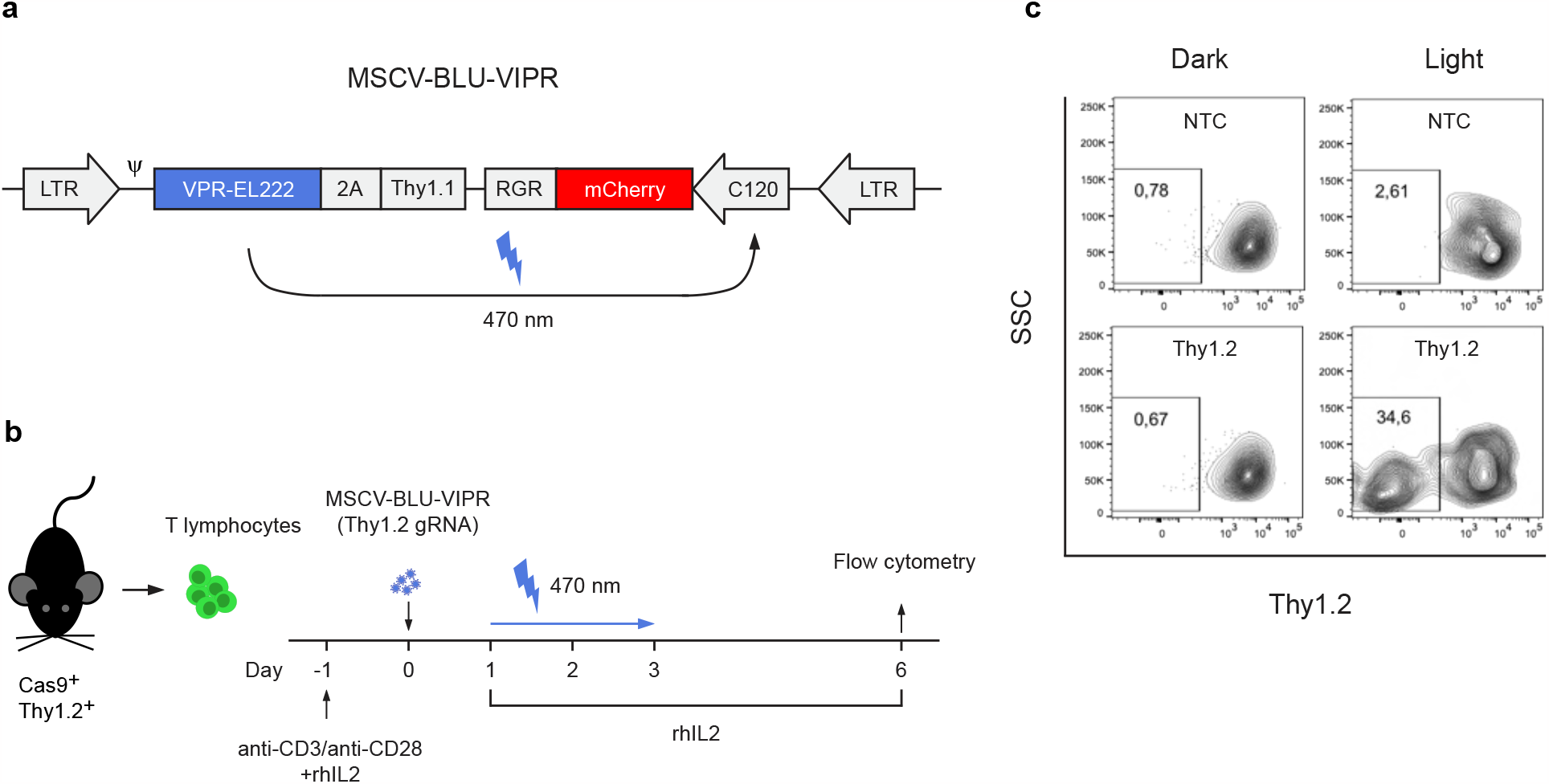
Optogenetic gene editing of primary mouse T lymphocytes in vitro. **a**, Design of the retroviral MSCV-BLU-VIPR constructs for blue-light induced expression of gRNA after transduction of primary mouse T lymphocytes. **b**, Cas9 transgenic splenic mouse T lymphocytes (Thy1.2^+^) where transduced with MSCV-BLU-VIPR containing Thy1.2 specific or non-targeting control (NTC) gRNA, followed by exposure to blue light and analysis by flow cytometry. **c**, After 48 hours of light exposure, followed by a 72 hour dark period, the T lymphocytes were stained for Thy1.2, gated for singlets and viability, and then analyzed for Thy1.2 expression. Representative result from two experiments is shown.

### Optogenetic gene editing of T lymphocytes in lymph nodes

Light-based gene perturbations of T lymphocytes in vivo would allow for spatiotemporal dissection of immune responses with unparalleled precision and pave the way for new insights into immune responses. This could, for example, be used to spatiotemporally dissect antitumoral immune responses to inform and improve immunotherapies against a variety of cancer types. To test if the BLU-VIPR system could be harnessed for precise optogenetic CRISPR in T lymphocytes in vivo, we first built an optogenetic setup allowing for precise illumination of individual lymph nodes in mice (Fig. 5a, b). We then asked if BLU-VIPR was tolerated by T lymphocytes in vivo, and to answer this we transduced Cas9 transgenic (CD45.1^+^) T lymphocytes with MSCV-BLU-VIPR followed by adoptive transfer into CD45.2^+^ TCRβ^-/-^ mice. Indeed, Cas9 transgenic T lymphocytes still expressed BLU-VIPR (Thy1.1^+^) after >20 weeks without loss of frequency or expression level, demonstrating that BLU-VIPR is tolerable to, and persists in, T lymphocytes in vivo (Extended Data Fig. 6). Next, we asked if we could achieve optogenetic gene editing in vivo, and to answer this we transduced Cas9 transgenic CD45.1^+^ T lymphocytes with MSCV-BLU-VIPR retrovirus containing NTC or Thy1.2 specific gRNAs followed by adoptive transfer into CD45.2^+^ TCRβ^-/-^ mice. Between three to six weeks after adoptive transfer, we delivered pulsed blue light (470 nm) to a single inguinal lymph node (iLN) to induce the expression of gRNA (Fig. 5c). After one hour of light exposure, we sutured the surgical incision, and 48 hours later we used flow cytometry to analyze the expression of Thy1.2 in transduced Cas9 transgenic T lymphocytes (CD45.1^+^ Thy1.1^+^) in secondary lymphoid organs and blood. To assess the efficiency of optogenetic gene editing we gated on transferred Cas9 transgenic T lymphocytes expressing BLU-VIPR (CD45.1^+^ Thy1.1^+^) and determined the frequencies of Thy1.2 positive (Thy1.2^pos^) and negative (Thy1.2^neg^) cells (Fig. 5d). Indeed, 48 hours after illumination of iLNs we found that >50% of CD45.1^+^ Thy1.1^+^ T lymphocytes transduced with Thy1.2 specific MSCV-BLU-VIPR virus were Thy1.2^neg^, demonstrating successful optogenetic gene editing in vivo (Fig. 5e). In contrast, Thy1.2 levels were not lost in CD45.1^+^ Thy1.1^+^ T lymphocytes in mice without exposure of iLNs to blue light, demonstrating that the MSCV-BLU-VIPR system requires blue light to induce gene editing (Fig. 5f). Cas9 transgenic T lymphocytes transduced with NTC control MSCV-BLU-VIPR virus did not lose expression of Thy1.2 after illumination (Fig. 5f). To further verify that the loss of Thy1.2 was induced by light illumination, we determined the expression of Thy1.2 on BLU-VIPR expressing Cas9 transgenic T lymphocytes (CD45.1^+^ Thy1.1^+^) cells in blood before and 48 hours after illumination of iLNs.

**Figure 5.**
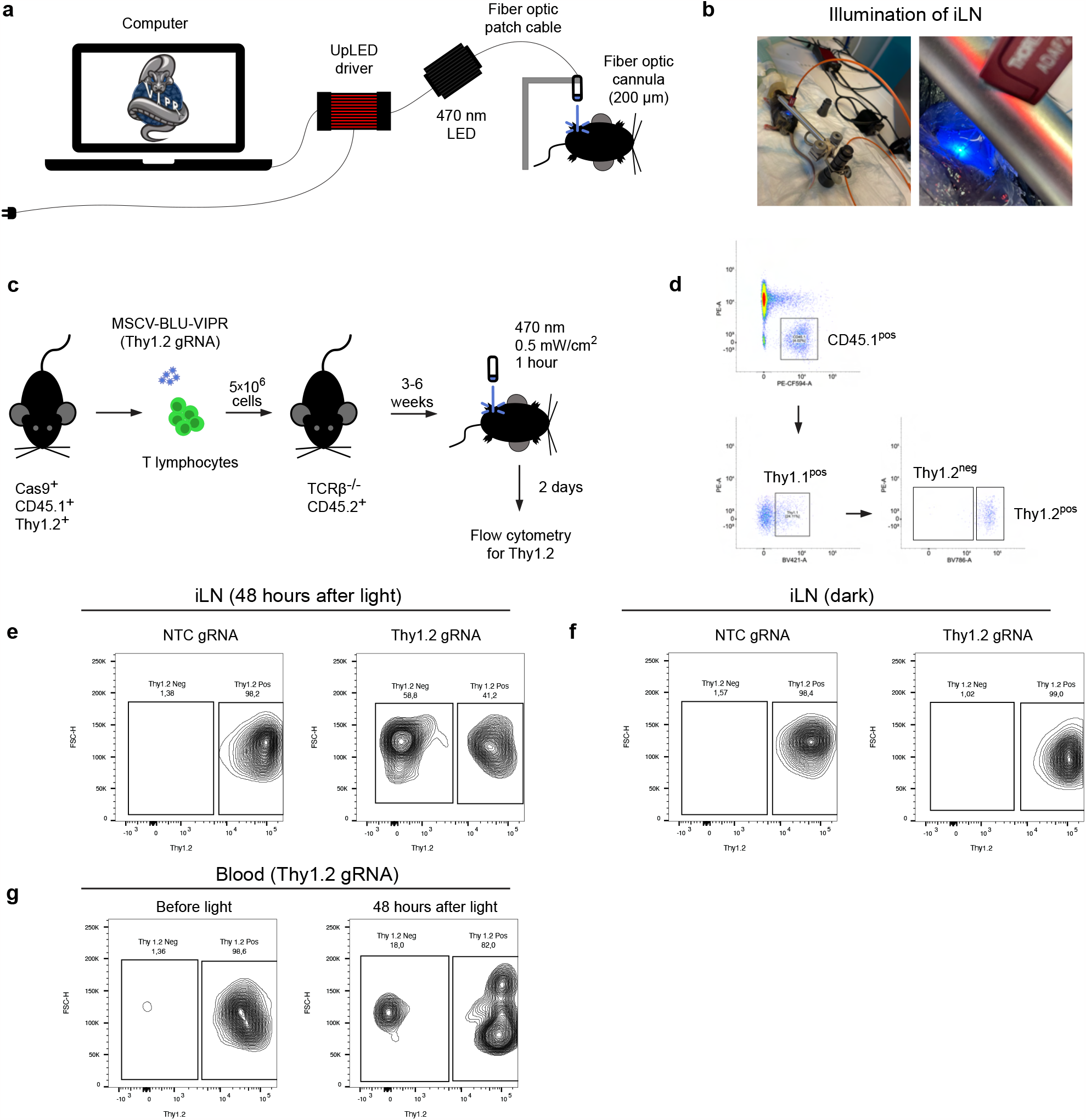
Optogenetic gene editing of Cas9 transgenic T lymphocytes in vivo. **a**, Optogenetic setup for illumination of lymph nodes with blue light (470 nm). **b**, Illumination of a single inguinal lymph node (iLN). **c**, Cas9 transgenic splenic mouse T lymphocytes (CD45.1^+^ Thy1.2^+^) where transduced with MSCV-BLU-VIPR containing Thy1.2 specific or non-targeting control (NTC) gRNA. The transduced cells were adoptively transferred to TCRβ^-/-^ CD45.2^+^ mice. After reconstitution of the T cell lymphocyte pool, we illuminated an iLN with blue light to induce gene editing of Thy1.2. **d**, Gating strategy to determine levels of Thy1.2 by flow cytometry. **e**, The levels of Thy1.2 on transduced Cas9 transgenic T lymphocytes (CD45.1^+^ Thy1.1^+^) from illuminated iLNs were determined by flow cytometry. **f**, The levels of Thy1.2 on transduced Cas9 transgenic T lymphocytes (CD45.1^+^ Thy1.1^+^) from non-illuminated mice were determined by flow cytometry. **g**, The levels of Thy1.2 on transduced Cas9 transgenic T lymphocytes (CD45.1^+^ Thy1.1^+^) in blood before and after illumination of an iLN were determined by flow cytometry.

Before illumination of iLNs no Thy1.2^neg^ CD45.1^+^ Thy1.1^+^ T lymphocytes were detected in blood, in contrast, after illumination >15% CD45.1^+^ Thy1.1^+^ T lymphocytes were Thy1.1^neg^ (Fig. 5g). In all, these results demonstrate that BLU-VIPR can be used for optogenetic gene editing in T lymphocytes in vivo.

## DISCUSSION

We have engineered a potent light-responsive transcription factor that can trigger the simultaneous expression of proteins and gRNAs from an RNA polymerase II promoter. The ribozyme-flanked gRNA design ensures precise excision of multiple gRNAs from a single transcript, therefore also allowing for multiplexed optogenetic gene editing. This new multipurpose optogenetic CRISPR platform can be combined with multiple Cas types and effectors and is deliverable by virus. Therefore, BLU-VIPR provides unprecedent flexibility for optogenetic CRISPR gene editing, allowing for efficient optogenetic knockouts in Cas9 transgenic primary T lymphocytes *in vivo*. The ability of BLU-VIPR to be introduced to primary cells using viral delivery systems confers several advantages, including its use in a wide range of cell types and its stable expression after integration in the genome. While the BLU-VIPR system is applicable to various cell types, tissues, and animal models, we chose to test the *in vivo* optogenetic gene editing in mouse T lymphocytes due to the clinical relevance of therapeutic T lymphocytes. Furthermore, since therapeutic T lymphocytes (e.g. tumor infiltrating lymphocytes and chimeric antigen receptor T lymphocytes) are generated *ex vivo* they could potentially be modified with BLU-VIPR before adoptive transfer to tumor-bearing recipients. This facile generation of optogenetic CRISPR murine T lymphocyte models could, therefore, be a useful tool for the precise spatiotemporal control of anti-tumor responses to improve tumor killing and reduce toxicities, in addition to allowing for the exploration of cancer immunotherapy questions requiring refined control of the genome in time and space.

While BLU-VIPR allows for unprecedented flexibility, as a gRNA-centered optogenetic platform, it is reliant on Cas being present in the target cell, however this can easily be achieved by co-delivery of Cas, or the use of Cas transgenic animal models. Likewise, the in vivo optogenetic model presented in this paper depends on homeostatic proliferation to reconstitute the T lymphocyte population in TCRβ knock out recipients. While reliance on homeostatic proliferation can be a drawback when investigating immune responses *in vivo*, adoptive transfer of MSCV-BLU-VIPR-transduced TCR transgenic, or Chimeric Antigen Receptor (CAR), T lymphocytes should circumvent this limitation^17–19^. Ultimately, BLU-VIPR enables the transformation of an off-the-shelf transgenic Cas model into an optogenetic gene perturbation-ready model. The usefulness of precise spatiotemporal perturbation of genes extends beyond immunology. Optogenetic techniques have been widely adopted by neuroscience, light being an excellent signal to control individual neurons. The understanding of the genetical underpinnings of neural circuitry and regulation of subsequent events could be elucidated with the help of optogenetic CRISPR tools. Similarly, multicellular organisms develop in highly regulated spatiotemporal patterns, making developmental biology and regenerative medicine natural candidates for optogenetic applications. In conclusion, the findings presented here could facilitate the introduction of efficient optogenetic CRISPR gene editing *in vivo* for use in experimental animal models across scientific fields.

## METHODS

### Animals

We used the following mouse strains: Cas9 (B6J.129(Cg)-Gt(ROSA)26Sor^tm1.1(CAG-cas9*,-EGFP)Fezh^/J), TCR ^-/-^ (B6.129P2-*Tcrb*^tm1Mom/J^), and CD45.1(B6.SJL-Ptprc^a^ Pepc^b^/BoyJ). All mice were housed in a specific pathogen-free animal facility at Center for Molecular Medicine, Karolinska Institutet. The study was approved by the Ethical Review Committee North, Stockholm County (Ethical approval Dnr 7547-2022), and animals were handled in compliance with the guidelines at Karolinska Institutet.

### Cell lines and viruses

HEK293T cells (laboratory stock), Platinum-E cells (CellBioLabs) and 293FT cells (Thermo Scientific) were cultured in high-glucose DMEM (Sigma Aldrich) supplemented with fetal calf serum (10%) (Sigma Aldrich), streptomycin (0.1 mg/ml) (Sigma Aldrich), penicillin (100 U/ml), Sodium pyruvate (1 mM) (Sigma Aldrich), HEPES (10 mM) (Sigma Aldrich) and L-glutamine (2 mM) (Sigma Aldrich). Cells were routinely tested for mycoplasma contamination. Transfections of HEK293T and 293FT cells were performed using X-tremeGENE 9 DNA Transfection Reagent (Roche). Murine Stem Cell Viruses (MSCV) were packaged using Platinum-E cells by co-transfection of MSCV retroviral transfer plasmids with pCL-Eco (Addgene #12371) using X-tremeGENE 9 DNA Transfection Reagent (Roche). Lentiviruses were packaged using 293FT cells by co-transfection of lentiviral transfer plasmids with pMD2.G (Addgene #12259) and psPAX2 (Addgene #12260) using X-tremeGENE 9 DNA Transfection Reagent (Roche).

### Reporter cell lines

Reporter cell lines for Cas9 and base editors were generated using lentiviruses made from CRISPR-SP-Cas9 reporter (Addgene #62733), pLV-SI-121 (Addgene #131126) and pLV-SI-112 (Addgene #131127). HEK293T cells were transduced in six-well plates (2×10^5^ cells/well) with CRISPR-SP-Cas9 reporter, pLV-SI-121 or pLV-SI-112 lentivirus followed by selection with puromycin (2.5 μg/ml).

### Gene synthesis, Gibson assembly, and Golden Gate assembly

We synthesized short (0.2-1 kb) gene fragments as GeneStrands (Eurofins) or gBlocks (Integrated DNA Technologies). Longer (1-5 kb) gene fragments were generated by gene synthesis (Eurofins). Alternatively, gene fragments were amplified from plasmids by PCR using Platinum Pfx DNA polymerase (ThermoFisher). Constructs were assembled by Gibson assembly using the Gibson Assembly® Master Mix (New England Biolabs) or by Golden Gate assembly. Assembled constructs were verified by Sanger sequencing and restriction digests. Other plasmids were obtained from commercial vendors or from the Addgene repository: pRL-CMV-Renilla (Promega), pRS0045 (Addgene #131124), pRS0035 (Addgene #131125), lentiCRISPR v2 (Addgene #52961), pLV-SI-121 (Addgene #131126), pLV-SI-112 (Addgene #131127), pCMV_ABEmax (Addgene #112095), pXR001 (Addgene #109049), pSBtet-GB (Addgene #60504), gRNA-Cloning vector (Addgene #41824), SP-dCas9-VPR (Addgene #63798), CRISPR-SP-Cas9 reporter (Addgene #62733), pCL-Eco (Addgene#12371), pMD2.G (Addgene #12259), and psPAX2 (Addgene #12260).

### Transilluminators

Two blue LED setups were designed and built for delivery of blue light to cells in culture. The first consists of a commercially available transilluminator (Large blue LED transilluminator, IO Rodeo), controlled by an Arduino microcontroller (UNO, Arduino) and a relay module (SRD-05VDC-SL-C, Songle). The transilluminator was mounted inside a cell culture incubator. The second setup was custom made for experiments with black walled, optical bottom 96 well plates. It consists of individually controlled LEDs (one LED per well), driven by LED drivers (STP16CP05, STM Microelectronics), directed by a TEENSY 3.2 microcontroller (PJRC). The LED array is housed in a standoff-supported, laser-cut ABS enclosure, aligning each LED to a determinate well. This system allows for individual control of light intensity and duration on a well-per-well basis. Both setups were calibrated for light intensity using a custom-built photodiode (PDB-C139, Advanced Photonics) Arduino pyranometer.

### Guide RNAs and primers

The gRNAs for Cas9 reporter, base editing reporters, *NEAT1* and *Thy1* were selected based on published gRNA sequences^12,16^.The NTC control gRNA was selected based on published gRNA sequences^20^. To identify potent gRNAs for CRISPRa we screened at least eight gRNAs for *BMP2* and *PDGFB* (not shown). The gRNA sequences used for the *BMP2* and *PDGFB* gRNAs were designed with the Genetic Perturbation Platform’s sgRNA Designer webtool (Broad Institute). gRNA sequences were chosen to target 75 to 300 base pairs upstream of the transcription start site using the Human GRCh38 reference genome, and to be used with SP-Cas9 (NGG PAM) or LB-Cas12a (TTTV PAM). All gRNAs and primers used in this study are listed on Extended Data Table 1 and 2.

### Quantitative RT-PCR

All RNA extractions were made using TRIzol (Invitrogen), and conversion of RNA to cDNA was performed using the High-Capacity cDNA Reverse Transcription Kit (Applied Biosystems). For gene expression analysis we performed RT-qPCR using TaqMan™ Universal Master Mix II, no UNG (Thermo Fisher Scientific) with the following TaqMan probes: *HPRT1* (Hs01003267_m1), *BMP2* (Hs00154192_m1), and *PDGFB* (Hs00966522_m1). A Light Cycler 96 real-time PCR system (Roche) was used for real-time PCR, and data were analyzed using the Light Cycler 1.1 software (Roche).

### Optogenetic knockouts and CRISPRa using BLU-VIPR

For the BLU-VIPR activity assays, 4×10^4^ cells HEK293T cells were seeded in black walled, optical bottom 96 well plates. Cells were transfected with 50 ng BLU-VIPR plasmid immediately after seeding. Twenty-four hours post-transfection, cells were exposed to 1 mW/cm^2^ of 470 nm light (Pulsed, 20 seconds ON, 60 seconds OFF) on a custom-made blue LED transilluminator for 24 hours. After light exposure, fluorescence micrographs were taken on a ZOE fluorescent cell imager (Bio-Rad). To measure optogenetic activation of Cas9, 4×10^4^ cells Cas9 reporter cells were seeded in black walled, optical bottom 96 well plates and transfected with 30 ng lentiCRISPRv2 (without gRNA) and 50 ng BLU-VIPR plasmids with targeting and non-targeting gRNAs. Twenty-four hours post-transfection, cells were exposed to pulses (20 seconds ON, 60 seconds OFF) of 1 mW/cm^2^ of 470 nm light or kept in the dark. After 48 hours of light exposure, fluorescence micrographs were taken on a ZOE Fluorescent Cell Imager (Bio-Rad). For CRISPRa assays, 4×10^4^ HEK293T cells were seeded in black walled, optical bottom 96 well plates and transfected with 50 ng SP-dCas9-VPR, or 3 ng LB-dCas12a-VPR, and 50 ng BLU-VIPR plasmids with targeting gRNAs. Twenty-four hours post-transfection, cells were exposed to pulses (20 seconds ON, 60 seconds OFF) of 1 mW/cm^2^ of 470 nm light, or kept in the dark, and harvested 24 hours later for RNA extraction and RT-qPCR.

### Optogenetic base editing using BLU-VIPR

Base editor reporter cells (4×10^4^ cells) were co-transfected with 50 ng pCMV_ABEmax or pRS0035 (Target-AID) with 50 ng of BLU-VIPR plasmids containing targeting or non-targeting gRNAs. After 24 hours, cells were exposed to pulses (20 seconds ON, 60 seconds OFF) of 1 mW/cm^2^ of 470 nm light or kept in the dark. For flow cytometric analysis of the EGFP reporter, cells were harvested 24 hours after light exposure and analyzed using a Gallios Flow Cytometer (Beckman Coulter). For imaging of the EGFP reporter, base editor cells were kept in the dark for 48 hours after light exposure and fluorescence micrographs were taken on a ZOE Fluorescent Cell Imager (Bio-Rad). To achieve optogenetic base editing of an endogenous gene, 4×10^4^ 293FT cells were seeded in black walled, optical bottom 96 well plates and transfected and transfected with 50 ng pRS0035 and 50 ng BLU-VIPR plasmids with *NEAT1* targeting or non-targeting gRNA. Twenty-four hours post-transfection, cells were exposed to pulses (20 seconds ON, 60 seconds OFF) of 1 mW/cm^2^ of 470 nm light for 48 hours or kept in the dark. Light-exposed cells were harvested and sorted for mCherry expression on a Sony SH800 cell sorter (Sony Biotechnology). Genomic DNA was extracted from cells exposed to light, and from cells kept in the dark, using the Monarch Genomic DNA Purification Kit following the manufacturer’s protocol (New England Biolabs). The targeted region in *NEAT1* was amplified by PCR (primers in Extended Data Table 2). Amplicons were sequenced by Sanger sequencing. Sequencing traces were analyzed with EditR software^14^.

### Retroviral transduction

Mouse T lymphocytes were isolated from Cas9 transgenic mice and transduced following a modified protocol by Kurachi et al^16^. Briefly, splenic T lymphocytes were isolated from Cas9 transgenic mice using negative selection with the EasySep Mouse T Cell Isolation Kit (Stemcell Technologies) and then activated with Dynabeads Mouse T-Activator CD3/CD28 beads (Cat # 11456D, ThermoScientific). The isolated T lymphocytes were cultured in optogenetic RPMI 1640 medium (Cat# 11835030, ThermoScientific), lacking phenol red and HEPES, supplemented with 200 U/ml recombinant human IL-2 (Peprotech). One day after activation, the T lymphocytes were transduced with MSCV-BLU-VIPR retrovirus by spinfection (2,000 rcf, 60 min). MSCV-BLU-VIPR retroviruses, containing Thy1.2 specific gRNA or non-targeting control gRNA, were packaged using Platinum-E cells.

### Optogenetic gene editing in primary mouse T lymphocytes

For optogenetic CRISPR in primary mouse T lymphocytes in vitro, Cas9 transgenic T lymphocytes were transduced with MSCV-BLU-VIPR in six-well plates and one day later the transduced cells were transferred to black-walled, optical bottom 96 well plates. T lymphocytes were then exposed for 48 hours to pulses of 1 mW/cm^2^ 470 nm light (20 seconds ON, 60 seconds OFF) or kept in dark conditions. Following this exposure period, all cells were kept in the dark for 72 hours and then stained for Thy1.2. The expression of Thy1.2 was analyzed using flow cytometry. For optogenetic CRISPR in vivo, Cas9 transgenic CD45.1^+^ T lymphocytes were transduced with MSCV-BLU-VIPR and one day later injected intravenously (5×10^6^ cells) into CD45.2^+^ TCR ^-/-^ mice. Following 3-8 weeks of homeostatic T cell expansion, the recipients were anesthesized (isoflurane 2%) and the mice were prepared for intravital optogenetic stimulation following a protocol adapted from Ulrich von Andrian.^21^ Briefly, the skin with the left inguinal lymph node was flipped inside out following a small incision immediately left to the midline and single suture traction was established to keep the LN exposed. The tissue was kept moist with isotonic saline (0.9% NaCl). A stimulation chamber was built around the lymph node using vacuum grease to prevent the isotonic saline from leaking out. The fiber optic cannula (Thorlabs) was placed directly over the lymph node. For analysis of optogenetic CRISPR experiments, secondary lymphoid organs and blood was collected 48 hours post-illumination and Thy1.2 expression was analyzed by flow cytometry.

### Flow cytometry

Dead cells were excluded by incubating with the dead cell stain SYTOX™ Red (Invitrogen) before acquisition according to the manufacturer’s protocol. To generate monostain controls for compensation we used Ultracomp Plus Beads (eBioscience), GFP BrightComp eBeads™ Compensation Bead Kit (ThermoScientific) and mCherry Flow Cytometer Calibration Beads (Takara). Flow cytometry of stained T lymphocytes was then performed using a BD LSRFortessa Cell Analyzer (BD Biosciences) and the data were analyzed using FlowJo v10 (BD). The following antibodies were used for flow cytometry: anti-mouse Thy1.2 Brilliant Violet 785 (clone 30-H12, BioLegend), anti-mouse Thy1.1 Brilliant Violet 421 (clone OX-7, BioLegend), anti-mouse CD45.1 PE/Dazzle 594 (clone A20, BioLegend), anti-mouse CD45.2-Alexa Fluor 532 (clone 104, Invitrogen (eBioscience)), anti-mouse CD3ε APC (clone 145-2C11, BD Pharmingen).

## ACKNOWLEDGEMENTS

We thank Dr. Sanjaykumar Boddul, Daniel Lake, Evangelos Doukoumopoulos and Roxana Reyna for valuable experimental contributions, and Dr. Fredrik Wermeling for kindly sharing Cas9 and CD45.1 mice. This study was funded by grants from National Institutes of Health (F31AR072502), Whitaker Foundation, Cancerfonden (20 0992 PjF), and Konung Gustaf V:s och Drottning Victorias Frimurarestiftelse.

## AUTHOR CONTRIBUTIONS

This study was conceptualized by DVP and AE. DVP: Designed all experiments, performed, or contributed to all experiments, analyzed all experiments, performed data analysis, and wrote the manuscript. NP: Contributed to mouse T lymphocyte experiments and to writing the manuscript. LF: Contributed to Cas12a CRISPRa and base editing experiments. TT: Contributed to CRISPRa experiments, transilluminator design and development, and to writing the manuscript. WN: Contributed to CRISPRa experiments and the design of the mouse T lymphocyte experiments.

GB: Contributed to writing the manuscript AE: Designed all experiments, analyzed all experiments, performed data analysis, and wrote manuscript.

