## Extended Data for "Light induced expression of gRNA allows for optogenetic gene editing of T lymphocytes in vivo"

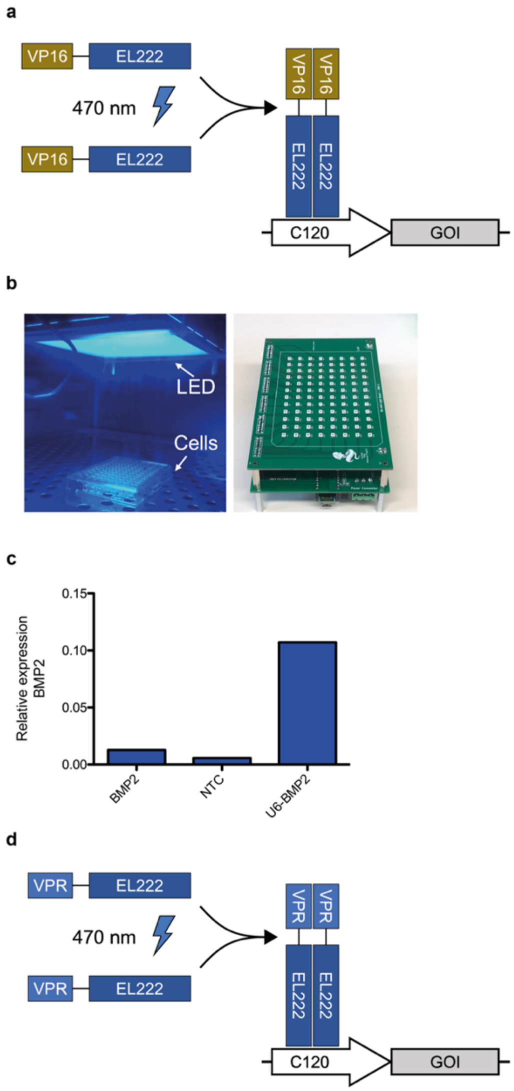


**Extended Data Figure 1. VP16-EL222 and VPR-EL222 Schematics and Initial test**
**a**, Schematics of the original VP16-EL222 system. **b**, Transilluminator blue LED setups used in the optogenetic experiments. Overhead transilluminator on the left, 96 well transilluminator on the right. **c**, VP16-EL222 based RGR production fails to activate significant expression of target endogenous gene with dCas9-VPR (U6-BMP2 is the same gRNA expressed constitutively from an hU6 promoter, positive control). Gene expression was normalized to *HPRT1* expression. The experiment was performed six times and one representative result is shown (mean of technical triplicates). **d**, Schematics of the VPR-EL222 system.

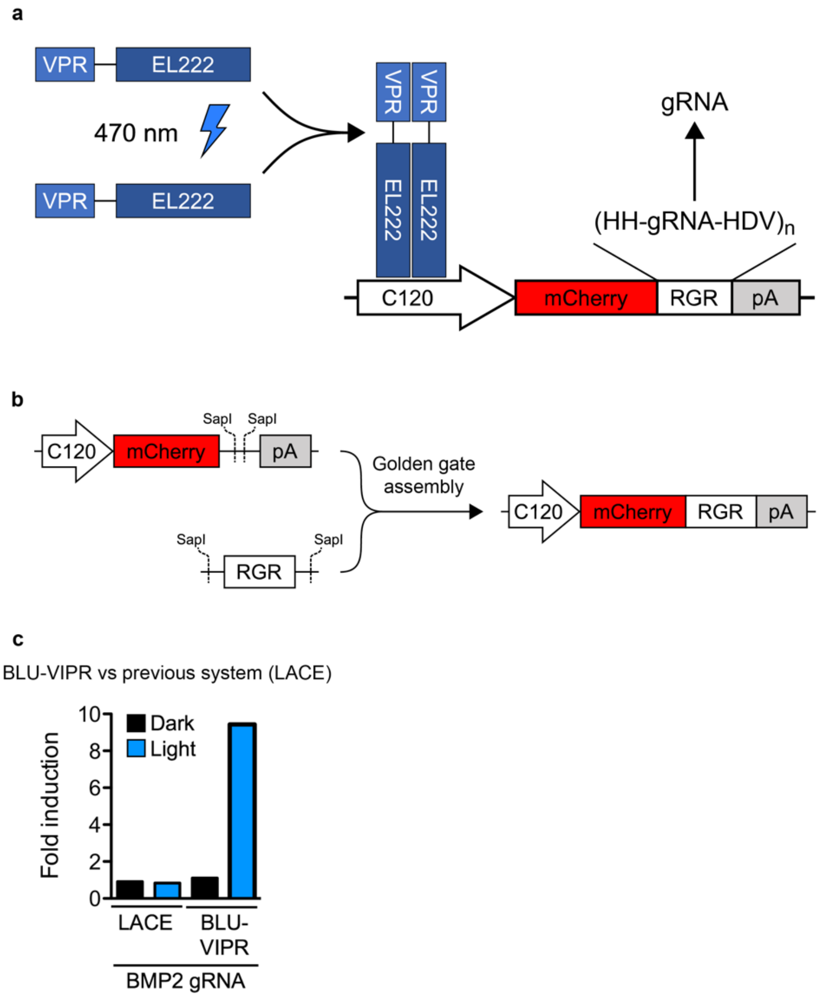


**Extended Data Figure 2. BLU-VIPR system schematics and Comparison to LACE**
**a**, Schematics of the BLU-VIPR system. **b**, Overview of cloning strategy for insertion of RGRs into BLU-VIPR plasmid. RGRs are designed with flanking SapI type IIS restriction enzyme sites, allowing for overhangs matching the vector backbone overhangs. A golden gate cloning, single step reaction results in the desired BLU-VIPR RGR plasmid. **c**, BLU-VIPR outperforms previously reported optogenetic CRISPRa system (LACE) when a single gRNA is used to target the endogenous *BMP2* gene. The experiment was performed three times and one representative result is shown (mean of technical triplicates).

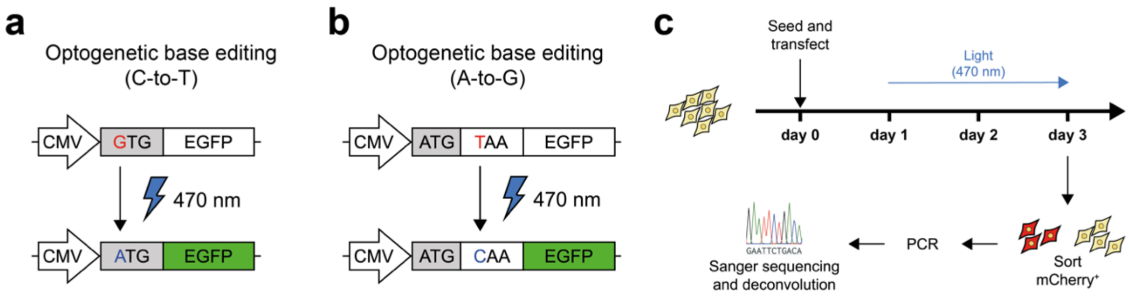


**Extended Data Figure 3. Optogenetic Base Editing Experimental Setups**
**a**, C-to-T base editing reporter cells turn EGFP positive after successful editing of C-to-T on the non-coding strand. **b**, A-to-G base editing reporter cells turn EGFP positive after successful editing of A-to-G on the non-coding strand. **c**, Overview of the optogenetic C-to-T base editing experiment targeting the endogenous lncRNA gene *NEAT1*.

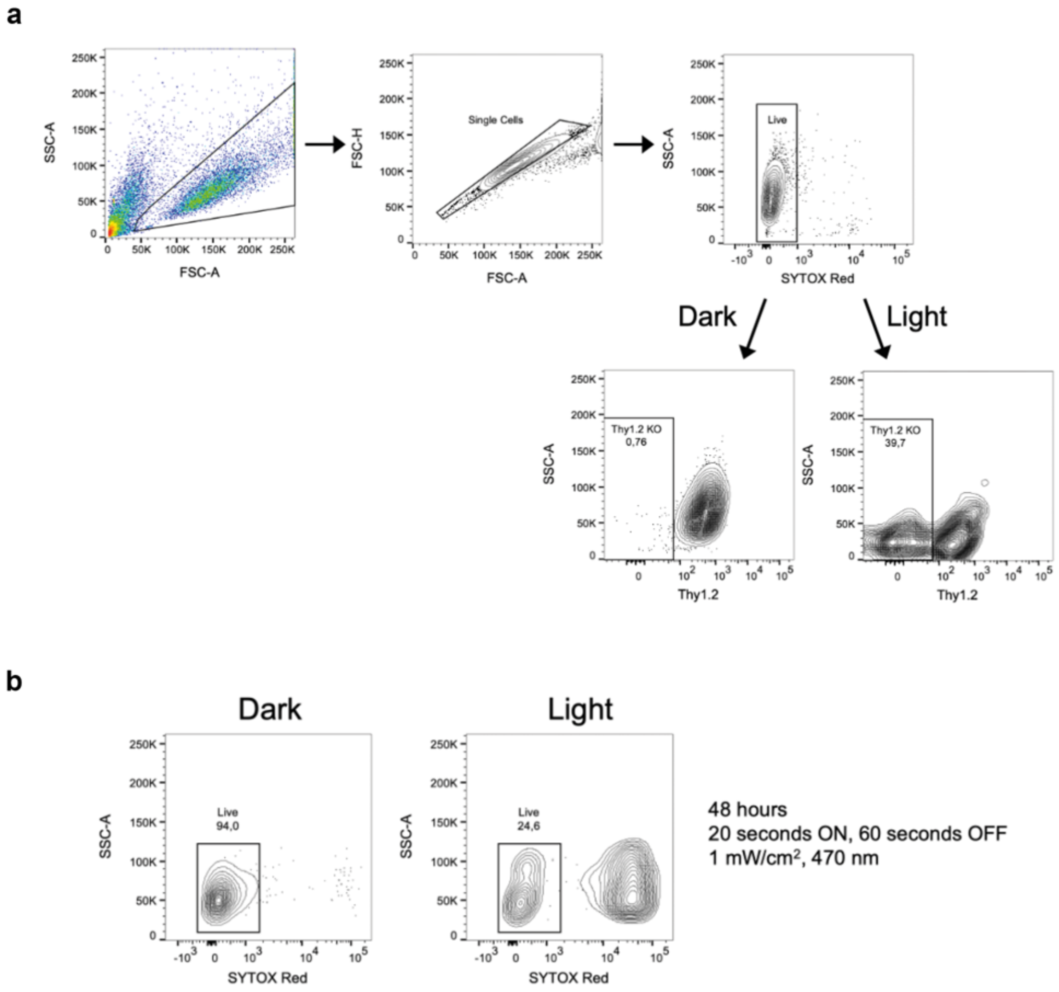


**Extended Data Figure 4. Gating Strategy for In Vitro T lymphocyte Experiments**
**a**, Gating strategy for optogenetic CRISPR experiment in Cas9 transgenic Thy1.2$^+$ mouse T lymphocytes. T lymphocytes were gated for single cells and viability (SYTOX$^{TM}$ Red negative) before being analyzed for Thy1.2 expression on the cell surface. **b**, The accumulation of toxic reactive species in vitro causes phototoxicity. Untransduced T lymphocytes where exposed to blue light for 48 hours and then stained with the dead cell stain SYTOX$^{TM}$ Red to estimate phototoxicity. SYTOX$^{TM}$ Red negative cells were considered viable. Representative result from two independent experiments is shown.

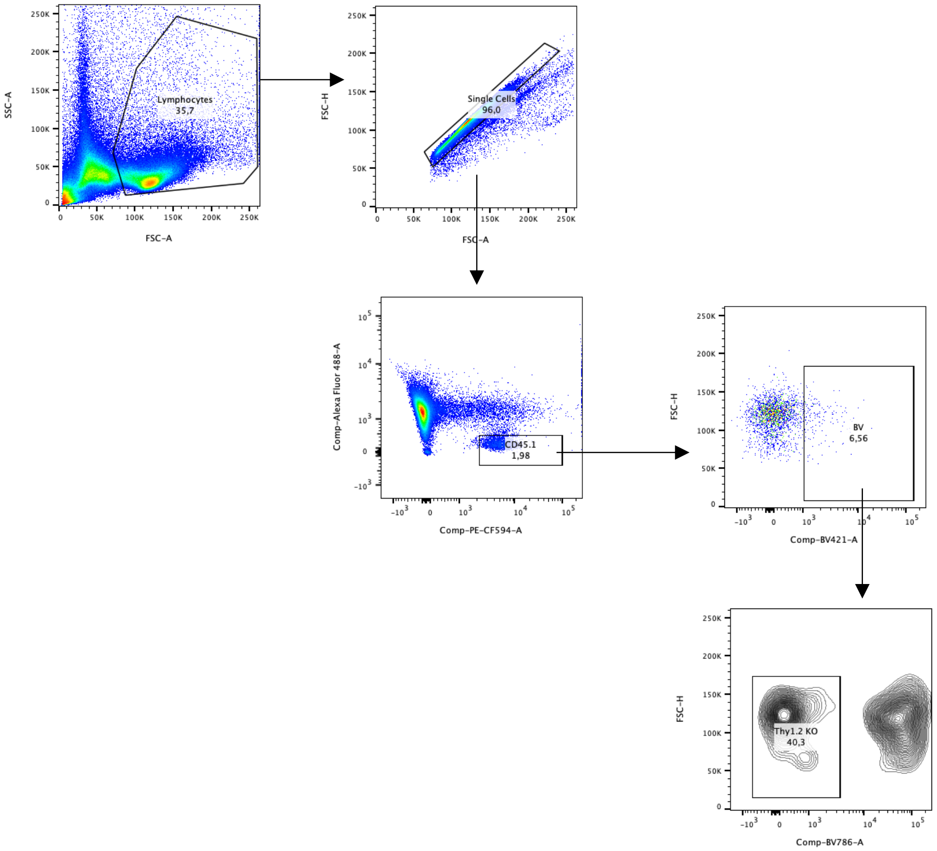


**Extended Data Figure 5. Gating Strategy for In Vitro T Lymphocyte Optogenetic Experiments**
Gating strategy for *in vivo* optogenetic CRISPR experiment in Cas9 transgenic CD45.1$^+$ mouse T lymphocytes. T lymphocytes were gated for single cells, CD45.1, and Thy1.1 (transduction marker) expression before being analyzed for Thy1.2 expression on the cell surface.

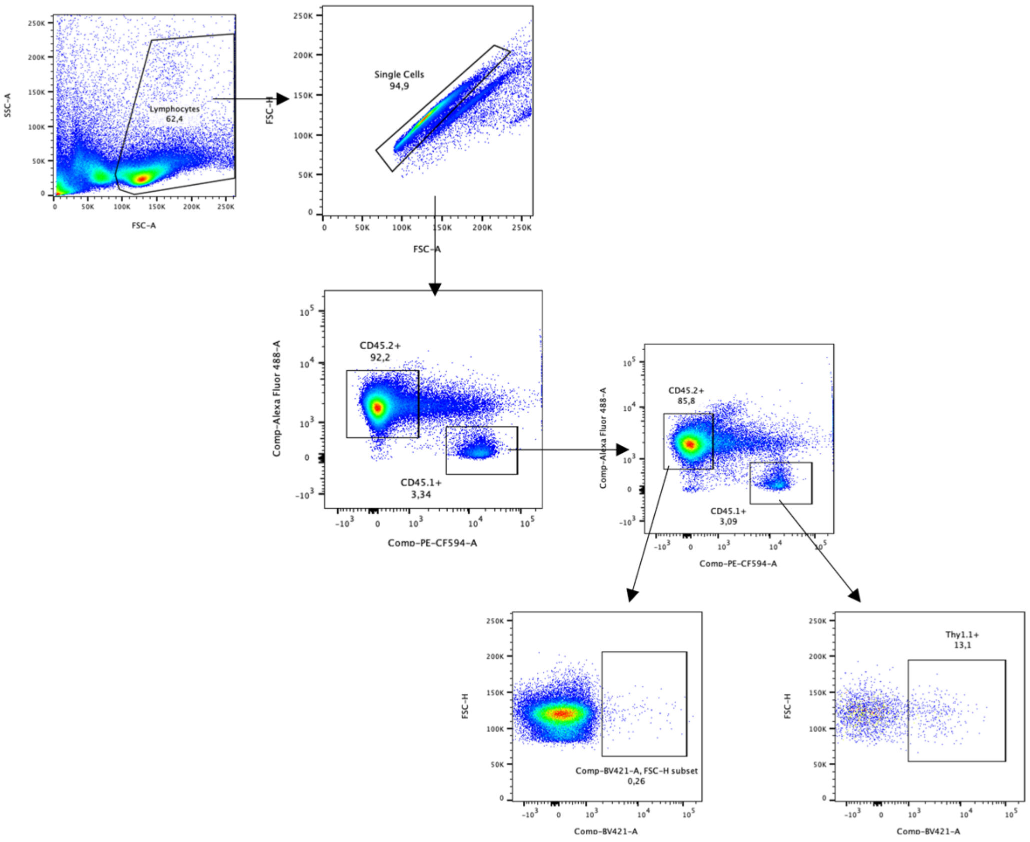


**Extended Data Figure 6. Gating Strategy for In Vivo T Lymphocyte Optogenetic Experiments**
Cas9 transgenic splenic mouse T lymphocytes ($CD45.1^+$, $Thy1.2^+$) were engineered *ex vivo* by transduction with MSCV-BLU-VIPR containing Thy1.2 specific or non-targeting control (NTC) gRNA. The transduced cells were adoptively transferred to $TCR\beta^{-/-}$ $CD45.2^+$ mice. After 20 weeks T lymphocytes were isolated from inguinal lymph nodes, and Thy1.1 expression was analyzed with flow cytometry.

**Extended Data Table 1**
List of all gRNA spacer sequences used in the optogenetic experiments.

| **Target** | **Spacer** | **Cas** |
|---|---|---|
| NTC (non-targeting control) | GAACGACTAGTTAGGCGTGTA | SP-Cas9 or LB-Cas12a |
| *BMP2* | GGCGAGCCGCGCCGCGAAGG | SP-Cas9 |
| Cas9 Reporter | GGGCCACTAGGGACAGGAT | SP-Cas9 |
| *PDGFB* | TAAAGGAGAAGGGAGAGTGCGAG | LB-Cas12a |
| C-to-T Reporter | CACGGTCACCCTGACACGCT | SP-Cas9 |
| A-to-G Reporter | CCTTATGACCCTGACACGCT | SP-Cas9 |
| *NEAT1* | GCCGACAGTGTCCAGTTCAG | SP-Cas9 |
| *Thy1* (Thy1.2) | CGTGTGCTCGGGTATCCCAA | SP-Cas9 |

**Extended Data Table 2**
Primers used for genomic DNA PCR amplification of *NEAT1* gene sequence surrounding the targeting window for the optogenetic base editing experiments.

| | |
|---|---|
| F | TAACTTACGGAGTCGCTCTACGGACTACCCCATCACAGAGTACTTTT |
| R | GGATGGGATTCTTTAGGTCCTGGGGTTTGTATGAACTTACTGGCATT |
